# Bacteria use processing body condensates to attenuate host translation during infection

**DOI:** 10.1101/2025.01.09.632196

**Authors:** Manuel González-Fuente, Nico Schulz, Alibek Abdrakhmanov, Gaiea Izzati, Shanshuo Zhu, Gautier Langin, Paul Gouguet, Mirita Franz-Wachtel, Boris Macek, Anders Hafrén, Yasin Dagdas, Suayib Üstün

## Abstract

Pathogens employ sophisticated strategies to modulate host protein homeostasis by targeting proteolytic pathways, but their impact on protein synthesis remains elusive. We report that pathogenic bacteria *Pseudomonas syringae* (*Pst*) targets ribonucleoprotein condensates, known as processing bodies (P-bodies), to attenuate host translation through two effectors with liquid-like properties. We uncovered a previously unknown link that *Pst*-mediated repression of the ER stress response is required for P-body assembly. Furthermore, we identify a novel intersection between P-bodies and autophagy, demonstrating that autophagic clearance of P-bodies is crucial for maintaining the balance between translationally active and inactive mRNAs. Altogether, our discoveries provide novel insights on how host translation is attenuated by bacteria to dampen plant immunity and uncover unknown connections between ER stress responses and autophagy with P-body dynamics.

---

Maintaining a proper balance between protein synthesis and degradation is crucial for all aspects of life. Protein homeostasis, or proteostasis, is key throughout all developmental stages and allows coordinated and timely responses to environmental stimuli and threats (*1*). As such proteostasis is targeted by pathogens in their efforts to modulate the host physiology to accommodate their own needs. Many pathogens interfere with plant proteolytic pathways such as the ubiquitin proteasome system or autophagy (*2, 3*). Similarly, biosynthetic pathways are also modulated by pathogens, particularly viruses, as expected considering their absolute reliance on the host translation machinery (*4, 5*). Other pathogens can also target protein synthesis pathways to promote disease: from modifying chromatin accessibility to directly hijacking transcription or mRNA maturation (*6*–*8*). However, despite the reported importance of translation for mounting adequate defense responses (*9, 10*), little is known about whether and how plant pathogens manipulate their host translation.

Defects in translation caused by internal cues or external stimuli cause the association of polysome-bound mRNAs to translation inhibitors (*11*). Translationally inactive mRNAs are then recruited by additional cytosolic proteins that mediate their phase separation into biomolecular condensates such as processing bodies (P-bodies) or stress granules, processes highly conserved among eukaryotes (*12*). In addition to the reversible sequestration of translationally inactive transcripts, P-bodies also harbour the mRNA decay machinery (*11*). P-bodies are involved in many developmental processes, abiotic stress responses and host-virus interactions in animals, fungi and plants (*13*–*15*). Moreover, P-bodies disassemble during the first layers of plant immunity (*16*). Nevertheless, to date it remains elusive whether P-bodies can be targeted by pathogens to modulate the host physiology.

Here we report that pathogenic bacterium *Pseudomonas syringae* pathovar *tomato* (*Pst*) deploys two effectors with liquid-like properties to induce the formation of plant P-bodies and promote disease. We revealed that *Pst* exploits plant P-bodies by repressing the endoplasmic reticulum (ER) stress response and hence attenuate translation, both processes important for mounting adequate defense responses (*9, 17*). Furthermore, we identified a novel intersection between P-bodies and autophagy, demonstrating that the autophagic clearance of P-bodies is crucial for maintaining the balance between translated and translationally arrested mRNAs. Altogether, our discoveries provide novel insights on how host translation is attenuated by bacteria via effector-triggered P-body formation to dampen plant immunity and uncover unknown connections between ER stress responses and autophagy with P-body dynamics.

## Results

### Pseudomonas syringae induces P-body formation in an effector-dependent manner

Considering the importance of biomolecular condensates in modulating translation, we sought to investigate the behaviour of P-bodies upon bacterial infection. We observed an increase in the number of P-body marker DECAPPING 1 (DCP1)-GFP foci 24 hours upon infection (Fig. 1A). Interestingly, this increase was compromised when using the type III secretion system-defective strain Δ*hrcC* (Fig 1B), indicating that P-body induction is effector-dependent. To rule out that this increase was caused by the ability of *Pst* to inhibit pathogen-associated molecular pattern (PAMP)-triggered immunity (PTI), we used the bacterial double Δ*avrPto/avrPtoB* mutant, which is unable to block the first stages of PTI signalling (*18*). The increase in DCP1-GFP foci number was not compromised upon infection with Δ*avrPto/avrPtoB* (Fig. S1A), indicating that inhibition of PTI is not the major cause of *Pst*-mediated increase in DCP1-GFP foci number. Similar behaviours were observed with additional GFP-tagged P-body components DCP5, VARICOSE (VCS) and SM-LIKE 1A (LSM1A; Fig. 1B and S1B-C). Moreover, both basal and *Pst*-induced DCP1-GFP and DCP5-GFP foci were sensitive to cycloheximide (CHX; Fig. S1D-E), suggesting that these condensates are *bona fide* P-bodies (*19*). Altogether, these results show that *Pst*, through its type III effectors, induces the formation of Arabidopsis P-bodies.

**Figure 1.**
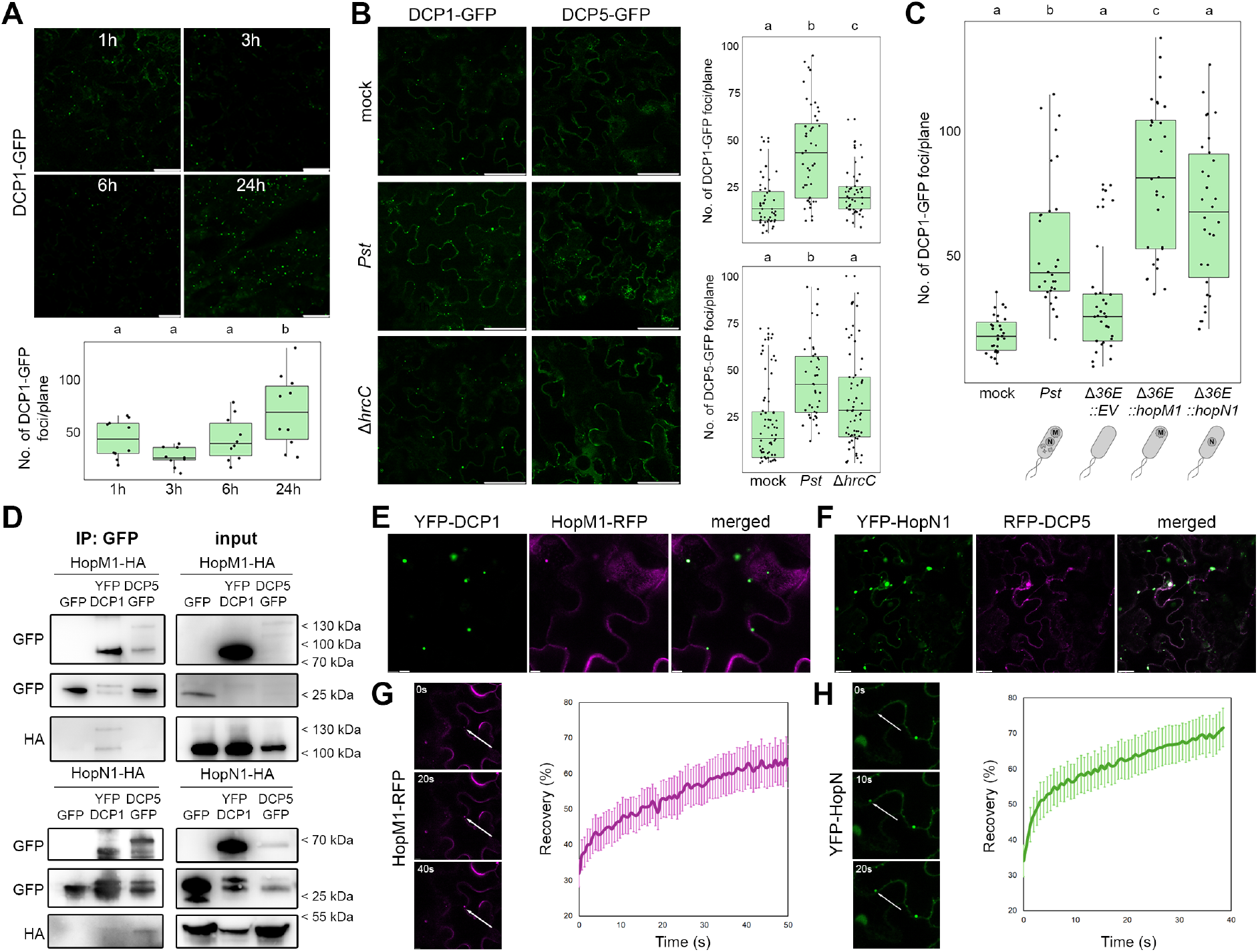
*Pseudomonas syringae* induces P-body formation in an effector-dependent manner. (**A**-**C**) Confocal microscopy pictures of epidermal cells of 4-week-old *Arabidopsis thaliana* leaves expressing P-body markers DECAPPING 1 (DCP1)-GFP (A-C) and DCP5-GFP (B) infected with 10^7^ CFUs/mL *Pseudomonas syringae* pathovar *tomato* DC3000 (*Pst*) wild-type or mutant strains, or 10 mM MgCl_2_ mock at different timepoints (A) or 24 hours (B-C) after mesophyll-infiltration. Accompanying boxplots represent the number of fluorescent foci (particles defined by an intensity threshold of 70-255) present per focal plane from at least two independent experiments. Scale bars depict 50 μm. Different letters indicate statistical groups determined using Student *t*-test (p-value < 0.05). (**D**) Coimmunoprecipitation of YFP-DCP1, DCP5-GFP or GFP with HA-tagged HopM1 or HopN1 transiently expressed in 5-week-old *Nicotiana benthamiana* leaves 24 hours after inoculation. Total proteins (input) were subjected to immunoprecipitation (IP) with GFP-Trap beads, followed by immunoblot analysis using either anti-GFP or anti-HA antibodies. GFP blot was split for visualization purposes. Representative result from two independent experiments. (**E-F**) Confocal microscopy pictures of transiently co-expressed YFP-DCP1 with HopM1-RFP (E) or YFP-HopN1 with RFP-DCP5 (F) in 5-week-old *N. benthamiana* epidermal cells. Images were taken 30–48 h after infiltration. Scale bars represent 5 μm. (**G-H**) Representative confocal microscopy images and time-course plot of the fluorescence signal recovery of HopM1-RFP (G) and YFP-HopN1 (H) foci after photobleaching. Recovery was quantified as the percentage of fluorescence intensity compared to before photobleaching. The data represented is the mean and error bars represent the standard error of the mean (n > 10). Scale bars represent 5 μm.

### HopM1 and HopN1 induce the formation and associate with P-bodies

To identify the effector(s) responsible for *Pst*-mediated increase in P-body number, ten single *Pst* effectors were transiently expressed in *Nicotiana benthamiana* to test their individual impact on YFP-DCP1 foci formation (Fig. S1F-G). Three effector proteins showed a significant increase in YFP-DCP1 foci number: Hrp outer protein M1 (HopM1), HopN1 and HopO1. For the first two, the P-body induction effect was further confirmed in a more natural infection of *A. thaliana* using singly delivered effectors through an effectorless *Pst* polymutant (*20*) (Fig. 1C). Additionally, in the case of HopM1, it had been previously detected as interacting with P-body components (*21*) (Fig. S1H), and P-body induction was partially compromised when infecting with a single Δ*hopM1* mutant (Fig. S1I). Altogether, these results point towards a specific role of certain effectors as inducers of P-body formation upon *Pst* infection.

To test whether the induction of P-body formation mediated by *Pst* effectors could occur in a direct or indirect way, we tested if P-body-inducing effectors HopM1 and HopN1 could associate with known P-body components. HA-tagged HopM1 co-immunoprecipitated with YFP-DCP1 and not with DCP5-GFP, whereas HopN1-HA showed the opposite (Fig. 1D). Similarly, YFP-DCP1 foci co-localized with HopM1-RFP signal, whereas DCP5-GFP foci did not (Fig. 1E and S1J). On the other hand, YFP-HopN1 foci seemed to partially co-localize with RFP-DCP5 and DCP1-RFP (Fig. 1F and S1K). Both HopM1-RFP and YFP-HopN1 presented liquid-like properties, as they both partially recovered their fluorescent signal upon photobleaching (Fig. 1G-H). Altogether, these results suggest that HopM1 and HopN1 associate specifically to different P-body components and are recruited to P-body-like structures likely through phase separation.

As HopN1 is a reported cysteine protease (*22*), we tested whether its P-body inducing ability was dependent on its enzymatic activity. Interestingly, a catalytic mutant version of HopN1, HopN1^D299A^ (*22*), was not able to induce YFP-DCP1 foci when transiently expressed in *N. benthamiana* (Fig. S1L-M), while partially retaining the ability to co-localize with P-bodies (Fig. S1N). Nevertheless, YFP-HopN1^D299A^ foci did not present liquid-like properties (Fig. S1O), indicating that its protease activity is indispensable for the recruitment of HopN1 to functional P-bodies and for the increase in their number. This strengthens the notion that prokaryotic effectors can undergo phase separation and that this is required for executing their proper mode of action (*23*).

### P-bodies are positive regulators of susceptibility via an altered endoplasmic reticulum stress response

Our results suggest that P-body formation might be a consequence of the bacterial manipulation of the host to accommodate *Pst* needs, rather than a plant defense response. To test this, we evaluated the bacterial multiplication in a P-body formation-defective mutant *dcp5-1* (*24*) (Fig. 2A). *dcp5-1* fostered a significantly lower bacterial multiplication than wild-type plants when naturally infecting with *Pst* (Fig. 2B). A similar pattern was observed when *Pst* was inoculated by mesophyll-infiltration, which bypasses the first layers of pre-invasive immunity (Fig. S2A). Furthermore, the canonical PTI defence response, flg22-induced reactive oxygen species (ROS) production, was not altered in *dcp5-1* (Fig. S2B). Altogether, these results suggest that P-bodies might be positive regulators of plant susceptibility and that the bacteria might actively induce their formation to their own benefit, beyond immunity suppression.

**Figure 2.**
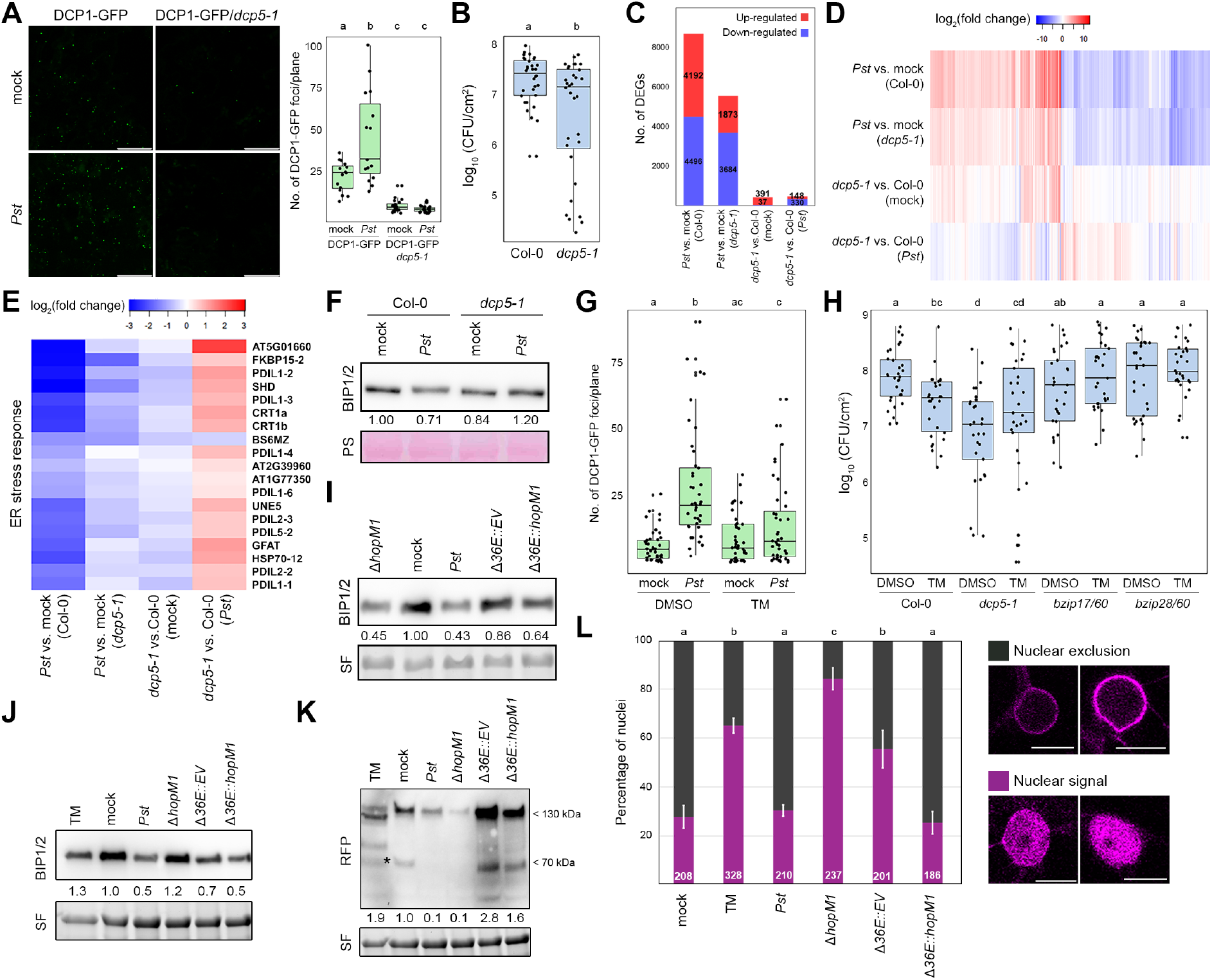
P-bodies are positive regulators of susceptibility via an altered ER stress response. (**A**) Confocal microscopy pictures of epidermal cells of 4-week-old *A. thaliana* leaves expressing DCP1-GFP in the wild-type Col-0 or mutant *dcp5-1* background 24 hours upon mesophyll-infiltration with 10^7^ CFUs/mL *Pst* or 10 mM MgCl_2_ mock. Accompanying boxplot represents the number of fluorescent foci present per focal plane from at least two independent experiments. Scale bars depict 50 μm. Different letters indicate statistical groups determined using Student *t*-test (p-value < 0.05). (**B**) Bacterial population three days post in 4-week-old *A. thaliana* leaves of wild type Col-0 and *dcp5-1* plants three days after dipping with 10^7^ CFUs/mL *Pst*. Data from three independent experiments. Different letters indicate statistical groups determined using Student *t*-test (p-value < 0.05). (**C**) Number of differentially expressed genes (DEGs) for each condition (i.e. treatment effects: *Pst* vs. mock-treated Col-0 and *dcp5-1* plants, and genotype effects: *dcp5-1* vs. Col-0 plants mock and Pst-treated). 4-week-old *dcp5-1* and Col-0 plants were mesophyll-infiltrated with 10 mM MgCl_2_ mock or 10^7^ CFUs/mL *Pst* and samples were harvested 24 hours after inoculation. Total number of DEGs per condition is depicted. Colours indicate the proportion of DEGs up-regulated (red) and down-regulated (blue). DEGs are considered when | log2(fold change) | > 1.5 and adjusted p-value < 0.05. (**D-E**) Heatmap representing the log_2_(fold change) of all DEGs defined in B (C) or a fraction of ER stress response-related genes (D). (**F**) Immunoblot analysis of ER stress response marker BINDING PROTEIN 1 and 2 (BIP1/2) present in 4-week-old *A. thaliana* leaves 24 hours after mesophyll-infiltration with either 10 mM MgCl_2_ mock or 10^7^ CFUs/mL *Pst*. The large subunit of the Rubisco visible after Ponceau S (PS) staining serves as loading control. Numbers indicate the band intensity of the BIP1/2 immunoblot normalized to each sample loading. Representative result from two independent experiments. (**G**) Number of fluorescent foci per plane observed in epidermal cells of 4-week-old *A. thaliana* leaves expressing DCP1-GFP infected with either 10 mM MgCl_2_ mock or 10^7^ CFUs/mL *Pst* and either 10 μM tunicamycin (TM) or equivalent Dimethyl sulfoxide (DMSO) volume, 24 and 8 hours prior imaging, respectively. Data from two independent experiments. Different letters indicate statistical groups determined using Student *t*-test (p-value < 0.05). (**H**) Bacterial population in 4-week-old *A. thaliana* leaves of wild type Col-0 and *dcp5-1* plants mesophyll-infiltrated with 10^4^ CFUs/mL *Pst* and either 6 μM TM or equivalent DMSO volume. Bacteria were extracted for counting three days post bacterial inoculation and two days post TM/DMSO treatment. Data from four independent experiments. Different letters indicate statistical groups determined using Student *t*-test (p-value < 0.05). (**I**) Immunoblot analysis of BIP1/2 present in 4-week-old *A. thaliana* leaves 24 hours after mesophyll-infiltration with either 10 mM MgCl_2_ mock or 10^7^ CFUs/mL of either wild-type *Pst*, single Δ*hopM1* mutant, the empty vector-complemented effectorless polymutant (Δ*36E::EV*) or the *hopM1*-complemented polymutant (Δ*36E::hopM1*). The large subunit of the Rubisco visible through Stain-Free imaging (BioRad) revelation serves as loading control. Numbers indicate the band intensity of the BIP1/2 immunoblot normalized to each sample loading. Representative result from two independent experiments. (**J-K**) Immunoblot analysis of BIP1/2 and RFP-BZIP28 present in 5-week-old *roq1 N. benthamiana* plants transiently expressing RFP-BZIP28 for 36 hours, and then mesophyll-infiltrated with either 10 mM MgCl_2_ mock or 10^7^ CFUs/mL of either wild-type or mutant *Pst* strains used in I, or 10 μM TM for 8 hours prior to sample harvesting. The large subunit of the Rubisco visible through Stain-Free imaging revelation serves as loading control. Numbers below the BIP1/2 immunoblot indicate the band intensity of the BIP1/2 immunoblot normalized to each sample loading. Numbers below the RFP immunoblot indicate the band intensity of the cleaved RFP-BZIP product, marked with and asterisk, normalized to each sample loading. Representative results from two independent experiments. (**L**) Percentage of imaged nuclei based on the RFP-BZIP28 signal in/around the nuclei of the samples depicted in J-K. Total number of nuclei per condition appears at the base of the barplot. The two categories were manually assigned as “nuclear excluded” (dark grey) or “nuclear imported” (magenta), as depicted in the accompanying figures, based on the intensity ratio between nuclear and perinuclear regions. The scale bar represents 10 μm. White error bars depict the standard error of the mean. Data from two independent experiments. Different letters indicate statistical groups determined using Student *t*-test (p-value < 0.05).

To further investigate the role of P-bodies during infection, we conducted transcriptomic analyses on *dcp5-1* and wild-type Col-0 upon infection (Table S1). Generally, the effect of the mutation was minimal compared to the effect of the bacterial infection, as the number of differentially expressed genes of *dcp5-1* in both mock and infected conditions was considerably small (Fig. 2C). Interestingly, when we compared the genes differentially regulated upon infection in both genotypes, we observed very similar tendencies (Fig. 2C-D and S2C). Therefore, it seems that P-bodies are not responsible for major changes during the transcriptional reprogramming occurring upon infection, including defense responsive genes (Fig. S2D). However, when carefully examining the differences between *dcp5-1* and Col-0 upon infection, we observed certain pathways were indeed distinctly regulated (Fig. S2C). Among the differentially down-regulated genes upon infection, we observed that ER stress responsive genes were strongly down-regulated in Col-0, but to a lesser extent in *dcp5-1* (Fig. 2E). This tendency was mirrored at the protein level using BINDING PROTEIN 1 and 2 (BIP1/2) as ER stress response markers (Fig. 2F). The ER stress response has emerged as a novel susceptibility hub (*17*), and ER exit sites have been recently linked to P-body organization in Drosophila (*25*). Therefore, we decided to further investigate the role of ER stress responses in modulating P-body dynamics upon infection.

In coherence with the differential regulation of ER stress responses in *dcp5-1* (Fig 2E-F), this mutant was found to be hypersensitive to ER stress-inducing drugs dithiothreitol (DTT) and tunicamycin (TM) when applied in seedlings (Fig. S2E). Moreover, TM had a negative effect on the number of YFP-DCP1 foci observed in seedling roots (Fig. S2F), evidencing a link between ER stress and P-bodies in plants. Indeed, chemical induction of ER stress in adult plants prevented Pst from inducing P-bodies (Fig. 2G and S2G). Additionally, TM had a deleterious effect in bacterial multiplication in Col-0, but not in *dcp5-1*, mirroring ER stress irresponsive double mutants *bzip17/60* and *bzip28/60* (Fig. 2H) (*26*). Altogether, these results point towards a deregulation of ER stress responses as a potential cause of the reduced susceptibility to *Pst* of *dcp5-1*, and evidence a tight link between P-bodies and ER stress responses in plants, as recently suggested in animals (*27*).

To further investigate the down-regulation of ER stress responses upon infection observed in Col-0 but absent in *dcp5-1* (Fig. 2E-F), and to evaluate the impact of *Pst* effectors therein, we blotted ER stress response markers BIP1/2 upon infection with the wild-type and effectorless polymutant *Pst* strains. As previously observed (Fig. 2F), *Pst* reduced the accumulation of BIP1/2, but the effectorless polymutant did not (Fig. 2I). This suggests a role for type III effectors in mediating the repression of ER stress responses. Analysing published transcriptomic data of the single Δ*hopM1 Pst* mutant (*28*), we observed that ER stress responsive genes were up-regulated (Fig. S2H), revealing a possible role for HopM1 as a suppressor of the host ER stress responses. Previous work indicated that HopM1 associates with trafficking components at the ER (*21*), strengthening our hypothesis that HopM1 may modulate ER stress responses. Indeed, we confirmed that *Pst* but not Δ*hopM1* repressed several ER stress responsive genes (Fig. S2I). Moreover, translocation of HopM1 alone was able to partially revert the BIP1/2 accumulation triggered by the effectorless polymutant (Fig. 2J). We then had a look at the status and subcellular localization of the ER-resident transcription factor BZIP28, which is processed and imported into the nucleus upon ER stress (*29*). Indeed, RFP-BZIP28 was cleaved and translocated to the nucleus upon TM treatment (Fig. 2K-L). Conversely, HopM1 reduced the processing and nuclear translocation of RFP-BZIP28 (Fig. 2K-L), supporting its role as major repressor of ER stress responses upon *Pst* infection and suggesting a possible molecular mechanism for this repression. Interestingly, the compromised induction of P-bodies by the Δ*hopM1* strain was not observed when plants were treated with TM (Fig. S2J), suggesting that ER stress repression is required for HopM1-mediated induction of P-bodies.

### P-bodies are required for Pst-mediated attenuation of the host translation

To understand the role and protein composition of P-bodies during infection, different proteomic approaches combining immunoprecipitation and proximity labelling coupled with mass spectrometry were conducted to determine the protein composition of *Pst*-induced P-bodies (Tables S2-3, Fig. S3A-E). Several known P-body components were identified validating our different proteomic datasets (Fig. S3A-B and S3E). Interestingly, the protein composition of *Pst*-induced P-bodies was quite different from the composition of basal P-bodies (Table S2), suggesting that these biomolecular condensates are not mere “kidnappers” of translationally inactive transcripts, but that they can also sequester proteins to modulate their function/accessibility in the cytosol (*30*). Several proteins never previously reported to associate with P-bodies were identified (Table S2). For their possible interconnections with protein homeostasis, among the newly identified P-body associated proteins, we confirmed the co-localization with P-body marker DCP1 and liquid-like properties of three: an uncharacterized protein hereafter referred as DCP1/5-INTERACTING PROTEIN 1 (DIP1; AT1G27752), the PROTEASOME SUBUNIT ALPHA TYPE 2-B (PAB2; AT1G79210) and the E3 ligase IAN9-ASSOCIATED PROTEIN 1 (IAP1; AT1G18660), recently described as RRS1-ASSOCIATED RING-TYPE E3 LIGASE (RARE; Fig. S3F-G) (*31, 32*).

Additionally, several proteins with links to translation were identified in our proteomics analyses (Table S3 And Fig. S1H and S3E). This is in agreement with the function of P-bodies as recruiters of translationally inactive transcripts in the cytosol (*11*). To test whether *Pst*-mediated induction of P-bodies had an impact on the host translation, we monitored translation activity using SUrface SEnsing of Translation (SUnSET) assays (*33*). Indeed, we observed that at the same timepoint at which *Pst* infection produced a major induction of P-bodies, there was a considerable reduction of newly synthesized polypeptides (Fig. 3A). This was compromised in *dcp5-1* (Fig. 3A), suggesting P-bodies modulate translation during infection. As an additional proxy for evaluating general translation, we monitored the phosphostatus of the RIBOSOMAL PROTEIN S6A (RPS6A). RPS6A is a downstream target of the master energy sensor TARGET OF RAPAMYCIN (TOR) involved in ribosome biogenesis and thus translation (*34*). Similarly to puromycin incorporation, RPS6A phosphorylation was severely reduced upon *Pst* infection, but to a lesser extent in *dcp5-1* (Fig. 3B). The observed *Pst*-mediated attenuation of translation is effector-dependent, as neither the single Δ*hrcC* mutant nor the effectorless polymutant dampened translation (Fig. 3C-D). This indicates that one or several effector(s) are responsible for *Pst* translational inhibition. P-body-inducing effectors HopM1 and HopN1 reverted the translation induction triggered by the effectorless polymutant, revealing their role as inhibitors of translation (Fig 3D). Interestingly, HopM1 was not able to repress translation in *dcp5-1*, indicating that HopM1 requires P-bodies to attenuate translation. On the other hand, HopN1 did repress translation both in *dcp5-1* and Col-0, decoupling P-body formation from translation for this particular effector and suggesting that these two effectors work differently. Altogether, these results highlight P-bodies as regulators of the *Pst*-mediated attenuation of translation.

**Figure 3.**
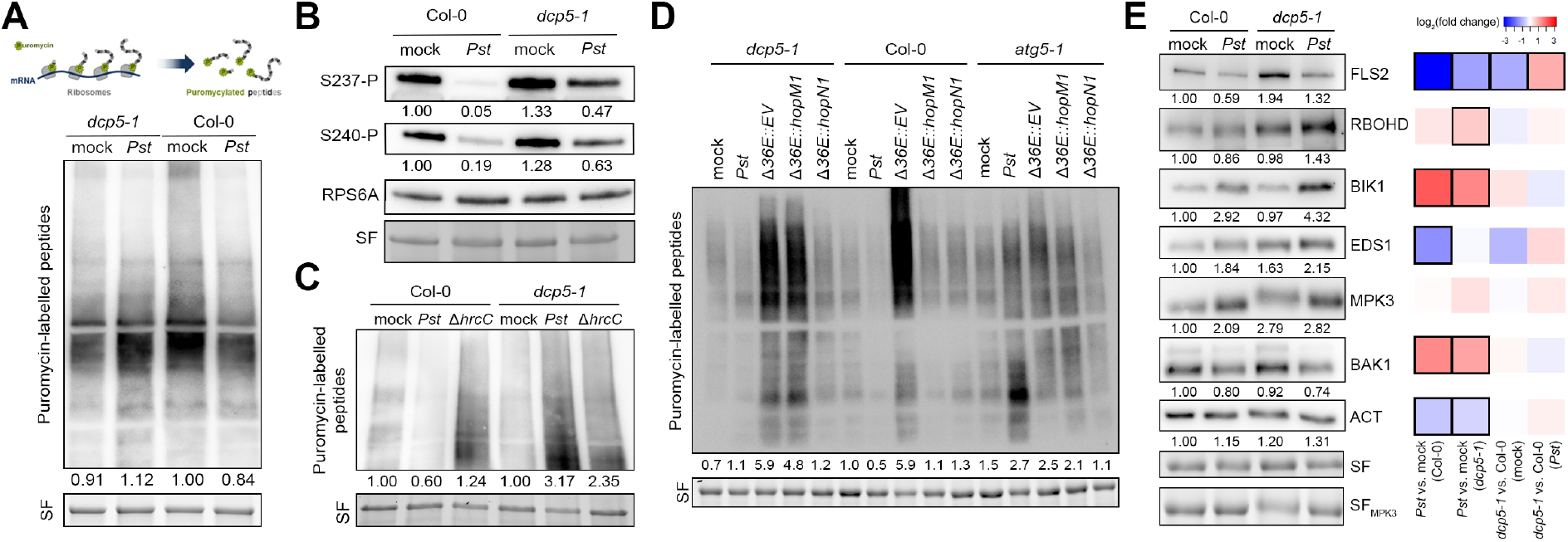
P-bodies are required for *Pst*-mediated attenuation of general translation. (**A** and **C-D**) Immunoblot analysis of puromycin-labelled peptides present in 4-week-old Col-0, *dcp5-1* (A and C), and *atg5-1* (D) *A. thaliana* leaves 24 hours after mesophyll-infiltration with either 10 mM MgCl_2_ mock or 10^7^ CFUs/mL wild-type or mutant *Pst* strains. The large subunit of the Rubisco visible through Stain-Free (SF) imaging revelation serves as loading control. Numbers indicate the lane intensity of the puromycin-labelled peptide immunoblot normalized to each sample loading. Representative results from at least two independent experiments. (**B**) Immunoblot analysis of the phosphorylation status at Ser237 and Ser240 of RIBOSOMAL PROTEIN S6A (RPS6A) in 4-week-old Col-0 and *dcp5-1 A. thaliana* leaves 24 hours after mesophyll-infiltration with either 10 mM MgCl_2_ mock or 10^7^ CFUs/mL *Pst*. The large subunit of the Rubisco visible through Stain-Free (SF) imaging revelation serves as loading control. Numbers indicate the band intensity ratios between the phosphorylation-specific RPS6A immunoblots and its respective phosphorylation-insensitive RPS6A blots. Representative result from two independent experiments. (**E**) Immunoblot analyses of immune components FLAGELLIN SENSITIVE2 (FLS2), RESPIRATORY BURST OXIDASE PROTEIN D (RBOHD), BOTRYTIS–INDUCED KINASE 1 (BIK1), ENHANCED DISEASE SUSCEPTIBILITY 1 (EDS1), MITOGEN-ACTIVATED PROTEIN KINASE 3 (MPK3) and BRI1-ASSOCIATED RECEPTOR KINASE (BAK1), and ACTIN (ACT) in 4-week-old Col-0 and *dcp5-1 A. thaliana* leaves 24 hours after mesophyll-infiltration with either 10 mM MgCl_2_ mock or 10^7^ CFUs/mL *Pst*. The large subunit of the Rubisco visible through Stain-Free (SF) imaging revelation serves as loading control. Numbers indicate the lane intensity of the puromycin-labelled peptide immunoblot normalized to each sample loading (two representative sets depicted in the figure). Representative results from at least two independent experiments. Accompanying heatmap depicts the log_2_(fold change) of the respective genes in the transcriptomic analyses from Fig. 2A. Black squares indicate that the corresponding adjusted p-value is lower than 0.05.

To investigate a possible pro-bacterial role of the P-body-mediated attenuation of host translation, we monitored the protein levels of several immune components. Five out of the six immune components studied (i.e. FLAGELLIN SENSITIVE2, FLS2; RESPIRATORY BURST OXIDASE PROTEIN D, RBOHD; BOTRYTIS–INDUCED KINASE 1, BIK1; ENHANCED DISEASE SUSCEPTIBILITY 1, EDS1; and MITOGEN-ACTIVATED PROTEIN KINASE 3, MPK3; but not BRI1-ASSOCIATED RECEPTOR KINASE, BAK1) presented higher protein levels in *dcp5-1* upon mock and/or *Pst* treatment, independently of their regulation at the transcript level (Fig. 3E). This shows that P-bodies are involved in the posttranscriptional regulation of gene expression of immune components (*35*), and suggests that *Pst* might attenuate plant translation as a virulence strategy to dampen defense responses.

### Pst utilizes autophagy to modulate P-body dynamics

Since *Pst* exploits HopM1 to suppress proteasome activity by inducing autophagy (*36*), we sought to investigate whether the effect of *Pst* on protein translation was dependent on these degradation pathways. We observed that *Pst*-mediated attenuation of translation was not majorly altered in the proteasomal mutant *rpt2a-2* (*37*), but it was completely deregulated in the autophagy-deficient mutant *atg5-1* (*38*) (Fig. 3D and S3H-I). To investigate the possible interplay between P-bodies and autophagy during infection we monitored P-bodies in *atg5-1*. We observed an increase both basally and upon *Pst* infection in the number of YFP-DCP1 foci (Fig. 4A), evidencing the role of autophagy in modulating P-body dynamics. This increase in YFP-DCP1 foci in *atg5-1* was accompanied by an increase in protein accumulation of YFP-DCP1 (Fig. 4B), suggesting that DCP1 might be a target of autophagic degradation. Indeed, DCP1 protein levels were decreased upon infection (Fig. 4B). This phenomenon was reverted when vacuolar degradation was chemically inhibited by concanamycin A treatment (Fig. S4A). Moreover, YFP-DCP1 co-localizes with mCherry-ATG8E-labelled autophagosomes in different autophagy-inducing conditions and (Fig. 4C and S4B), supporting the notion that DCP1 is a *bona fide* target of autophagic degradation. However, additional P-body components DCP5, LSM1A or VCS were not visibly degraded upon *Pst* infection, although DCP5 protein levels were basally stabilized in *atg5-1* (Fig. S4C). Interestingly, the YFP-DCP1 foci observed in *atg5-1*, both in mock and infected conditions, were also considerably larger in size (Fig. 4A) and insensitive to the translation inhibitor CHX (Fig. 4D and S4D). This suggests that, although increased in number, these P-bodies might not be completely functional or might not sequester translationally inactive transcripts. Additional lines of evidence toward this hypothesis include that these enlarged P-bodies are not sensitive to neither *Pst*-mediated induction nor TM-mediated repression (Fig. 4E and S4E). This is also in agreement with our previous findings that translation is not attenuated but rather induced in *atg5-1* (Fig. 3D and S3H). Interestingly, the P-body- and autophagy-inducing effector HopM1 seems to require autophagy for its translational repression activity (Fig. 3D). Altogether, these results point towards a complex interplay between P-bodies and autophagy, which is required for both *Pst*-mediated induction of P-bodies and subsequent translation attenuation, and for their proper functioning and turnover.

**Figure 4.**
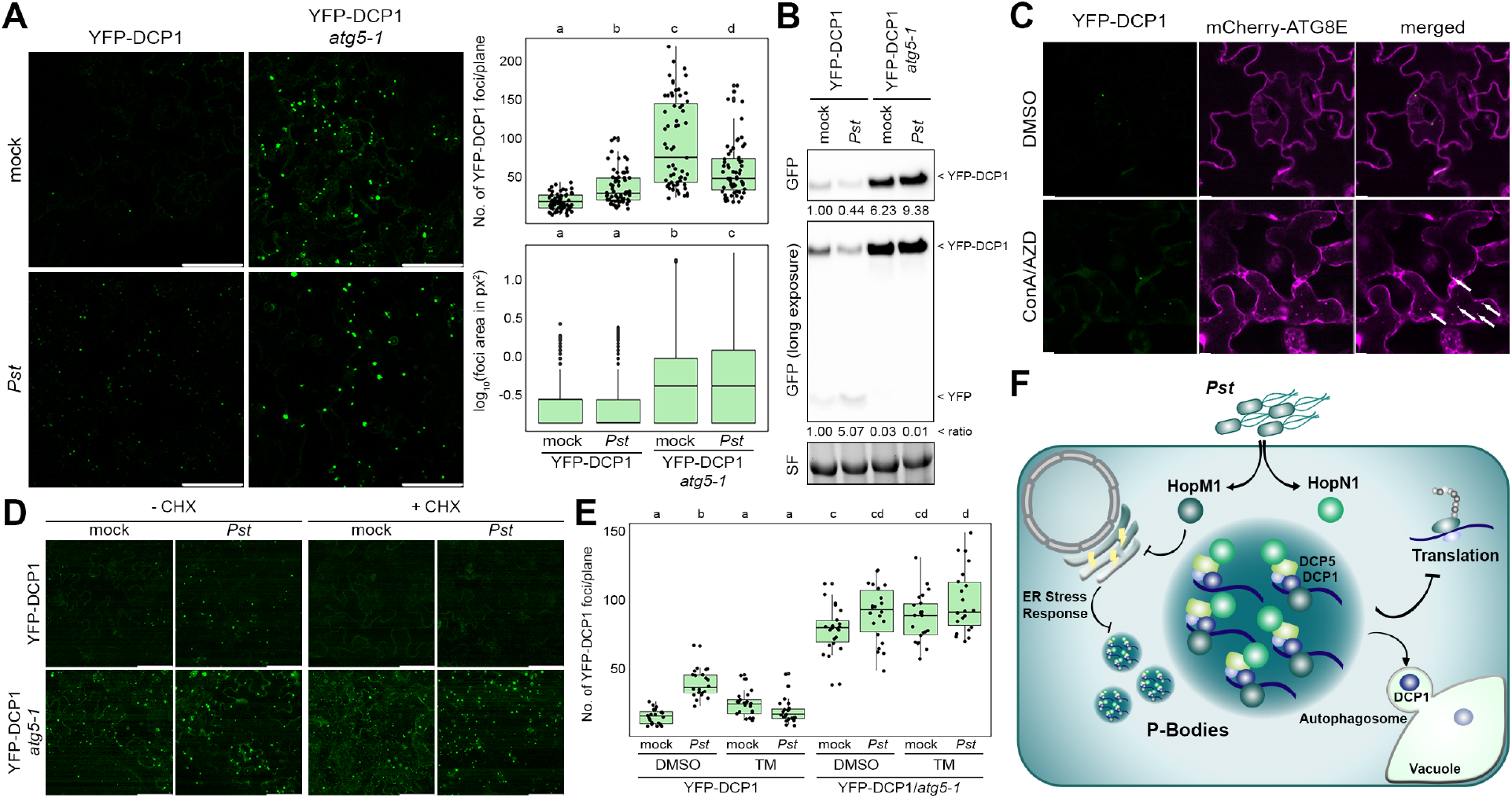
*Pst* utilizes autophagy to modulate P-body dynamics. (**A**) Confocal microscopy pictures and boxplots representing the number and size of fluorescent foci per plane observed in epidermal cells of 4-week-old *A. thaliana* leaves expressing YFP-DCP1 in the Col-0 and *atg5-1* backgrounds 24 hours after mesophyll-infiltration with 10^7^ CFUs/mL Pst or 10 mM MgCl_2_ mock. Scale bars depict 50 μm. Data from two independent experiments. Different letters indicate statistical groups determined using Student *t*-test (p-value < 0.05). (**B**) Immunoblot analysis of YFP-tagged protein levels in 4-week-old *A. thaliana* leaves expressing YFP-DCP1 in the Col-0 and *atg5-1* backgrounds 24 hours after mesophyll-infiltration with 10 mM MgCl_2_ mock or 10^7^ CFUs/mL *Pst*. The large subunit of the Rubisco visible through Stain-Free (SF) imaging revelation serves as loading control. Number indicate the band intensity of YFP-DCP1 normalized to each sample loading (top) and the ratio of YFP/YFP-DCP1 band intensity (bottom, depicted as “ratio”). (**C**) Confocal microscopy pictures of epidermal cells of 4-week-old *A. thaliana* leaves expressing YFP-DCP1 and mCherry-ATG8E 24 hours after mesophyll-infiltration with 10 mM MgCl_2_ mock or 10^7^ CFUs/mL *Pst*. Scale bars depict 5 μM. White arrows highlight co-localization of YFP-DCP1 and mCherry-ATG8E foci. (**D**) Confocal microscopy pictures of epidermal cells of 4-week-old *A. thaliana* leaves expressing YFP-DCP1 in Col-0 or *atg5-1* backgrounds mesophyll-infiltrated with 10 mM MgCl_2_ mock or 10^7^ CFUs/mL *Pst* 24 hours before imaging, and vacuum-infiltrated with 20 μM CHX or equivalent water volume 4 hours before imaging. Scale bars represent 50 μm. (**E**) Number of fluorescent foci per plane observed in epidermal cells of 4-week-old *A. thaliana* leaves expressing YFP-DCP1 in the Col-0 and *atg5-1* background mesophyll-infiltrated with either 10 mM MgCl_2_ mock or 10^7^ CFUs/mL *Pst* 24 hours before imaging, and 6 μM TM or equivalent DMSO volume 8 hours before imaging. Data from two independent experiments. Different letters indicate statistical groups determined using Student *t*-test (p-value < 0.05). (**F**) Model for *Pst*-mediated induction of P-body formation and attenuation of translation: *Pst* translocates effectors HopM1 and HopN1 that associate with host P-body components DCP1 and DCP5 respectively, to induce P-body assembly and attenuate protein translation. Additionally, *Pst* represses the host ER stress responses and induces autophagy, both processes required for induction of P-bodies and translation attenuation.

Since not all P-body components seem to be degraded via autophagy (Fig. S4C), suggesting substrate specificity, we sought to investigate whether some of the newly identified P-body-associated proteins (Fig. S3A-B and Table S2), could be also degraded via autophagy. Among the studied candidates (Fig. S3F), we selected DIP1 as it contains a Coupling of Ubiquitin conjugation to ER degradation (CUE) domain. Since this domain is known to bind ubiquitin (*39*), a common signal for autophagic degradation (*1*), we explored whether DIP1 could be involved in the interplay between autophagy and P-bodies. We confirmed that DIP1 co-localizes and co-immunoprecipitates with P-body components DCP1 (Fig. S3F and S4F). Moreover, similarly to DCP1, DIP1 is also degraded by autophagy as its protein levels are stabilized upon inhibition of vacuolar degradation or autophagy by concanamycin A treatment and co-expression of AIM peptide (*40*), respectively (Fig. S4G-H). Altogether, these results show that DIP1, similarly to DCP1, is also a target of autophagic degradation, suggesting that DIP1 might be involved in the interplay between autophagy and P-body homeostasis.

## Discussion

Pathogens have evolved sophisticated strategies to manipulate the host proteostasis, particularly targeting the proteasome and autophagy pathways (*2, 3*). However, how pathogens manipulate the other half of the proteostasis balance, notably at the translation level, is poorly understood. Here we show that *Pst* drives an enhanced condensation of P-bodies to attenuate protein translation via two effectors with liquid-like properties. Moreover, we showed that *Pst*-mediated repression of ER stress response and induction of autophagy is required for P-body assembly and translation attenuation (Fig. 4F).

The ability to induce phase separation is emerging as a strategy employed by pathogens to form biomolecular condensates (*41*). Recent studies indicate that pathogenic bacteria effectors exhibit high propensity for phase separation and drive condensation (*23*), although their exact mechanisms remain unclear. We shed light into how effectors with such properties might induce the assembly of P-bodies and provide the first evidence that bacteria can exploit this to attenuate the host translation to the pathogen own benefit. While we show that P-body induction leads to translational arrest, we cannot exclude that their condensation may also sequester proteins to render them nonfunctional or to neo-functionalize them, as recently suggested (*42*). Given that protein translation is fine-tuned upon pathogen perception (*9, 10*), it is unsurprising that pathogens have evolved strategies to manipulate this process by targeting P-bodies. Our findings also imply that other mechanisms are utilized by *Pst* to dampen host translation (e.g. TOR activity inhibition). But what advantage does *Pst* gain from dampening host translation? Certain immune components are known to be more actively translated during plant immune responses (*35*). Our analysis suggests that P-bodies play a crucial role in regulating the synthesis of these proteins. As such, future research will focus on other strategies to target host translation and on identifying specific mRNAs affected in translation efficiency, potentially uncovering whether they include immune components, susceptibility hubs or other pathways critical to limit the infection.

The molecular mechanisms behind P-body assembly are less understood. For decades, researchers have debated the factors driving P-body formation, which involves the aggregation of smaller ribonucleoprotein particles, protein-RNA interactions, low-complexity protein sequences, and liquid-liquid phase separation (*11*). While some P-body components are required for their maintenance, it is unclear how they are formed or their implications in pathologically relevant stress conditions (*14*). Our findings show that *Pst* triggers P-body formation in an effector-dependent manner and highlight novel mechanistic insights into how ER stress responses control P-body assembly. Considering the association between P-bodies and the ER (*27*), as well as the role of ER contact and exit sites in regulating P-body organization (*25*), targeting the ER and dampening ER stress responses appear to be efficient strategies for inducing P-body assembly. Additionally, we provide evidence that *Pst*-mediated induced autophagy regulates P-body assembly and disassembly, as analogously observed in studies on Kaposi’s sarcoma-associated herpesvirus (*43*). Emerging evidence highlights the roles of the ubiquitin-proteasome system and autophagy in controlling the assembly and disassembly of biomolecular condensates like stress granules and P-bodies (*44, 45*). These findings underscore the significance of protein degradation pathways as regulators of stress-responsive biomolecular condensates.

But why does *Pst* degrade P-bodies? We speculate that the proper function of P-bodies requires their recycling. This is supported by our observation that autophagy deficiency makes P-bodies resistant to CHX treatment, which generally leads to a reduction of untranslated mRNAs (*46*). P-bodies typically form only in the presence of untranslated mRNAs (*11*). As such our findings strongly suggest that without recycling, P-bodies cannot effectively sequester mRNAs. Alternatively, it is possible that selective autophagy of P-bodies is necessary to eliminate translationally inactive mRNAs, preventing them from re-entering new cycles of translation.

Collectively, our study reveals that *Pst* manipulates protein translation by inducing P-body condensation via the concerted action of at least two bacterial effectors. Utilizing these effectors as cellular probes, we uncover a previously unknown connection between the ER stress response and P-bodies, identifying the pathways and factors required for P-body formation. Considering that protein translation and P-bodies are conserved across kingdoms, our findings suggest that similar mechanisms may operate in other host-microbe interactions.

## Acknowledgments

We thank David S. Guttman, Darrell Desveaux (University of Toronto), and Thomas Lahaye (ZMBP) for the effectorless polymutant and complemented *Pst* strains, and Damien Garcia (IBMP, CNRS/University of Strasbourg) for the *dcp1-3, dcp5-1*, YFP-DCP1, DCP1-GFP, DCP5-GFP *A. thaliana* lines and plasmids. We thank Lea Röhder (RUB), Linus Börnke (Heinrich Heine University of Düsseldorf), Tom Denyer (ZMBP), Silke Wahl and Irina Droste-Borel (Proteome Center Tübingen) for technical support.

## Author contributions

Conceptualization: M.G.F., S.Ü.

Investigation: M.G.F., S.Ü., N.S., S.Z., G.L., G.I., P.G.

Visualization: M.G.F., S.Ü.

Resources: A.A., A.H., Y.D.

Funding acquisition: M.G.F., S.Ü.

Project administration: M.G.F., S.Ü.

Supervision: M.G.F., B.M., S.Ü.

Writing – original draft: M.G.F., S.Ü.

Writing – review & editing: M.G.F., S.Ü.

## Data and materials availability

The mass spectrometry data from this publication will be made available on the PRIDE archive, with the identifiers PXDXXXXX and PXDXXXX. All relevant transcriptomic and proteomic data are available in the supplemental information.

## Supplementary Materials

Materials and Methods

Figs. S1 to S4

Tables S1 to S8

