## Supporting information S1-4 for "Bacteria use processing body condensates to attenuate host translation during infection"

### Materials and Methods

Plant material and growth conditions

Arabidopsis thaliana ecotype Col-0 wild-type and its derived T-DNA mutants and transgenic plants listed in Table S4 were grown under either short day or long day conditions (light/dark cycles: 16h 22°C/8h 20°C, 130 $\mu mol \cdot mm^2 \cdot s^{-1}$ light intensity, 70% relative humidity) for *in vitro* experiments and seed propagation, or short day conditions (light/dark cycles: 12h 22°C/ 12h 20°C, 90 $\mu mol \cdot mm^2 \cdot s^{-1}$ light intensity, 70% relative humidity) for bacterial infection assays. For *in vitro* experiments, seeds were sterilized with 70% ethanol and sown on solid 1% sucrose-supplemented ½ Murashige and Skoog (MS) medium. For experiments with adult plants, seeds were sown on individual 6-cm-diameter pots filled with soil and, for infection assays, plants were grown for 4-5 weeks. Seeds for both experiments *in vitro* and with adult plants were stratified for 2 days at 4ºC. Nicotiana benthamiana plants were grown under long day conditions (light/dark cycles: 16h/8h, at 21°C and 70% humidity) in 6-cm-diameter individual pots filled with perlite-supplemented soil for 4-6 weeks.

Bacterial strains and growth conditions

*Pseudomonas syringae* pathovar *tomato* wild type strain DC3000 and its derived mutants listed in Table S5 were grown in King's B (KB) medium supplemented with 100 mg/L rifampicin (plus 25 mg/L kanamycin and 50 mg/L spectinomycin for the Δ*36E* strains) at 28ºC. *Agrobacterium tumefaciens* C58C1 derived strains listed in Table S5 were grown in Luria-Bertani broth (LB) medium supplemented with the appropriate antibiotics (i.e. 150 mg/L kanamycin, 300 mg/L gentamycin, 100 mg/L spectinomycin, 5 mg/L tetracycline and 50 mg/L rifampicin) at 28ºC. Escherichia coli TOP10 and DB3.1 derived strains were grown in LB medium supplemented with the appropriate antibiotics (i.e. 150 mg/L kanamycin, 300 mg/L gentamycin, 100 mg/L spectinomycin, 25 mg/L chloramphenicol and 50 mg/L carbenicillin).

Molecular cloning

The list of plasmids used and generated in this study are listed in Table S6. The list of primers used to amplify the coding sequences from *A. thaliana* genes are listed in Table S7. For Gateway® cloning, attB1 and attB2-flanked proof-reading PCR products were introduced into entry plasmids using BP clonaseTM II enzyme mix (Thermo Fisher Scientific). Expression vectors were generated using LR clonaseTM II enzyme mix (Thermo Fisher Scientific). Chemically competent E. coli TOP10 cells were transformed by heat shock (45 seconds at 42ºC, 2 min at 4ºC and 1h recovery at 37ºC prior to plating on antibiotic-supplemented plate). Positive clones were verified by colony PCR and Sanger sequencing (Eurofins).

Plant inoculation and bacterial growth assays

Overnight *Pst* cultures were centrifuged for 7 minutes at 4,000 rpm. Pellets were washed twice with 10 mM $MgCl_2$ prior to measuring the optical density at 600 nm ($OD_{600}$). For P-body quantification and transcriptomic, proteomic and gene expression analyses, *Pst* inocula were prepared at a final $OD_{600}$ of 0.1 in 10 mM $MgCl_2$. For bacterial growth assays, the inocula were prepared at a final $OD_{600}$ of 0.0001 in 10 mM $MgCl_2$. The bacterial inocula were infiltrated with a needleless syringe into the abaxial side of fully developed leaves from 4-5-week-old *A. thaliana* plants. For bacterial growth assays upon dipping inoculation, the final $OD_{600}$ was adjusted to 0.1 in 10 mM $MgCl_2$ and 0.025% Silwet L-77. Plants were fully dipped in the bacterial solution for 1 minute. After inoculation, all plants were well-watered and put in covered trays to reach nearly saturated humidity conditions. Unless stated otherwise, plants were used for experiments 24 hours after inoculation for P-body quantification and transcriptomic, proteomic and gene expression analyses, or 3 days post inoculation for bacterial growth assays. Bacterial populations were estimated by plating serial dilutions of bacteria extracted from two 6-mm-diameter leaf discs in 200 µL 10 mM $MgCl_2$ homogenized with a tissue lyser (Retsch). Colony forming units were manually counted 48 hours after plating.

Transient expression in Nicotiana benthamiana leaves

Overnight *A. tumefaciens* cultures were centrifuged for 7 min at 4,000 rpm. Pellets were washed twice with agroinfiltration buffer (10mM $MgCl_2$, 10mM MES pH 5.7) prior to measuring $OD_{600}$. Inocula were prepared at a final $OD_{600}$ of 0.1-0.5 in 200µM acetosyringone-supplemented agroinfiltration buffer and incubated at room temperature and darkness for 2 hours before infiltration with a needleless syringe into the abaxial side of 4 to 6-week-old *N. benthamiana* leaves. Plants were used for further experiments 24-48 hours after infiltration.

Laser scanning confocal microscopy and image analysis

Images were taken with a confocal laser scanning microscope (STELLARIS 8; Leica) using a water-immersed objective (HC PL APO CS2 63×/1.20 W) with a resolution acquisition set to 512 x 512 (P-body quantification and FRAP analyses) or 1024 × 1024 (for subcellular localization analyses) and a line average of 2-3. For GFP, excitation was set to 488 nm and emission was recorded at 500–560 nm. For YFP, excitation was set to 514 nm and emission was recorded at 530–570 nm. For RFP and mCherry, excitation was set to 587 nm and emission was recorded at 600–640 nm. Chlorophyll autofluorescence was visualized in the far-red wavelength. After acquisition, images were processed using Application Suite X (LASX; Leica) and ImageJ. For clarity purposes, contrast on the individual channels was manually adjusted. For P-body quantification (number and size), the analyze particle tool of ImageJ was used, counting for particles whose intensity threshold were between 70 and 255 on images taken with identical acquisition setup. Fluorescence recovery after photobleaching (FRAP) assays were conducted using the FRAP-wizard from the Application Suite X (LASX; Leica). The recovery overtime of independent photobleached foci was recorded and averaged before plotting. For RFP-BZIP28 nuclear translocation analyses, several images were taken with an identical acquisition setup. Afterwards, the signal around/inside every imaged nucleus was manually assigned as either excluded from the nucleus or imported into the nucleus as depicted in Fig. 2L.

Immunoblotting

Proteins were extracted in extraction buffer (100 mM Tris pH 7.5, 1 mM EDTA, 3% SDS) and mixed with 4X Laemmli buffer (BioRad) prior to boiling at 95ºC for 10 minutes and clearing by centrifugation of 1 minute at 13,000 rpm. Total protein extracts were separated by SDS–PAGE, visualized with the Stain-Free Imaging Technology in a ChemiDoc Imaging System (BioRad), transferred to PVDF membranes (BioRad), blocked with 5% skimmed milk in TBS (200 mM Tris and 1500 mM NaCl), for 1 hour at room temperature and incubated with the correspondent antibody listed in Table S8 for 1-2 hours at room temperature or overnight at 4ºC. For the detection, the immunoreaction was developed using an ECL Prime Kit (GE Healthcare) and detected with a ChemiDoc Imaging System (BioRad). When the Stain-Free images of the total protein extracts were not taken, Ponceau S staining (0.1% Ponceau Red and 5% acetic acid) was performed on the membranes after revelation instead. The relative quantification of the protein band intensities and ratios was performed using the Image Lab software (BioRad).

Co-immunoprecipitation

Plant tissue was grinded with a mortar and liquid nitrogen and then extracted in 1ml/g of GTEN buffer (10% glycerol, 25 mM Tris pH 7.5, 1 mM EDTA, 150 mM NaCl, 1 mM DTT, 1X Protease inhibitor Cocktail (Merck) and 0.5% Triton X,). The solution was vortexed until homogenized and incubated at 4ºC with rotation for 10min following centrifugation for 30 minutes at 4000 g and 4°C. Supernatant was filtered with three layers of Miracloth (Merck). 50µl of the filtrate was sampled as input, supplemented with 12.5µL Laemmli buffer 4X (BioRad) and boiled for 10 minutes at 95°C. Next, 10µl/g of GFP-Trap® Agarose beads (ChromoTek) were added to the remaining filtrate and incubated for 2 hours at 4ºC. The filtrates were then centrifuged at 800 g for 1 minute, the supernatants were carefully removed, and the pelleted beads were transferred to new 1.5mL microcentrifuge tubes in 1mL of GTEN buffer. Five steps of washing were performed by centrifuging at 800 g for 1 minute using 1mL GTEN buffer. After the last washing, 2X Laemmli buffer (BioRad) was added to equal amounts of beads and the samples were boiled 10min at 95°C prior to immunoblotting as previously described.

Measurement of ROS production

16 4-mm-diameter leaf discs were harvested for each condition and individually incubated overnight in a 96-well plate (PierceTM; Thermo Scientific) containing 100 µL of distilled water. For ROS production measurements, water was replaced by 100 µL of elicitor mix (50 nM flg22, 100 µM luminol, 20 µg/mL horseradish peroxidase) or mock (elicitor mix without flg22) and relative luminescence was quantified in each well every minute during 1 h, using a plate reader (Tecan). The total amount of luminescence per condition was calculated as the average of the sums of all measurements taken during the recorder hour per sample.

RNA extraction and quantitative real-time PCR

Total RNA from plant samples was extracted following the manufacturer instructions with either the RNeasy Plant Mini Kit (Qiagen) for RNA sequencing, or with the NucleoZOL reagent (Macherey-Nagel) for the rest of applications. After extraction, 500 ng of total RNA per samples were treated with DNase I (Thermo Scientific) prior to the cDNA synthesis using the LunaScript® RT SuperMix Kit (New England Biolabs), in both cases following the manufacturer instructions. Quantitative real-time PCR was performed using 2X MESA Blue qPCR Master Mix (Eurogentec) using a 2-step protocol for 40 cycles and melting curve analysis. The relative gene expression was then calculated following the ΔΔCt method. The primers used for qPCR are listed in Table S7.

RNA sequencing

Sequencing libraries were prepared from the extracted RNA using the NEBNext® UltraTM II RNA Library Prep Kit for Illumina® and Dual Index Primers (New England Biolabs) following the manufacturer instructions. The quality of the libraries was verified with a Bioanalyzer (Agilent) prior to send for Illumina® sequencing at the NovaSeq PE150 Platform of Novagene (Cambridge, UK). After quality control, reads were mapped to the A. thaliana TAIR10 reference genome. The mapped reads were counted with htseq-count for subsequent analysis. The $\log_2$ of the fold change (log2FC) and false discovery rate (FDR) were determined using R package DEseq2 with default settings (*47*). A gene is considered significantly differentially expressed when the absolute value of its log2FC > 1.5 and its FDR < 0.05.

IP-MS/MS and proximity labelling

IP-MS/MS analyses were conducted as previously described (*48*). Biotinylation test and proximity labelling assays were also conducted as previously described (*49*).

For purification, proteins were subjected to a NuPAGE 12% gel (Invitrogen) and in-gel trypsin digestion was done on Coomassie-stained gel pieces with a modification: *chloroacetamide* was used instead of iodoacetamide for carbamidomethylation of cysteine residues to avoid formation of lysine modifications isobaric to two glycine residues left on ubiquitinylated lysine after tryptic digestion. Next, peptides mixture were desalted using C18 Stage tips and run on an Easy-nLC 1200 system coupled to a Q Exactive HF mass spectrometer (both Thermo Fisher Scientific) as described (*50*) with some modifications: separation of the peptide mixtures was done using a 76-min segmented gradient from 10-33-50% of HPLC solvent B (80% acetonitrile in 0.1% formic acid) in HPLC solvent A (0.1% formic acid) at a flow rate of 200 nl/min. The seven most intense precursor ions were sequentially fragmented in each scan cycle using higher energy collisional dissociation (HCD) fragmentation. In all measurements, sequenced precursor masses were excluded from further selection for 30 s. The target values were $10^5$ charges for MS/MS fragmentation and $3 \times 10^6$ charges for the MS scan.

Acquired MS spectra were processed with MaxQuant software package version 1.5.2.8 with an integrated Andromeda search engine. Database search was either performed against an *Arabidopsis thaliana* database (41,609 protein entries, downloaded on 7th of October 2020 (*51*), and the sequences of DCP1-GFP and DCP5-GFP, or against *Nicotiana benthamiana* (74,802 protein entries (*51*), and the sequences from *A. thaliana* DCP1, *Escherichia coli* GUS and *Pst* HopM1. In both processes 285 commonly observed contaminants were included, too.

Endoprotease trypsin was defined as a protease with a maximum of two missed cleavages. Oxidation of methionine, phosphorylation of serine, threonine, and tyrosine, GlyGly dipeptide on lysine residues, and N-terminal acetylation were specified as variable modifications. Carbamidomethylation on cysteine was set as a fixed modification. Peptide mass tolerance was set to 4.5 parts per million (ppm) for precursor ions and 20 ppm for fragment ions. Peptide, protein, and modification site identifications were reported at a false discovery rate (FDR) of 0.01, estimated by the target-decoy approach (Elias and Gygi). The iBAQ (Intensity Based Absolute Quantification) and LFQ (Label-Free Quantification) algorithms were enabled, as was the "match between runs" option (*52*).

For the downstream analysis, $\log_2$(fold change) and FDR for each protein group were determined as described previously (*48*, *53*).In the case of the proximity labelling proteome analysis, conduced in *N. benthamiana*, to each protein group a unique Agi-code was assigned by blasting every *N. benthamiana* protein present in the protein group against *A. thaliana* proteome and taking the most frequent Agi-code with the best E-value. An E-value threshold of $e^{-10}$ was used, in case of absence of *A. thaliana* the protein group was not considered for downstream analysis.

<u>SUrface SEnsing of Translation (SUnSET) assays</u>

SUnSET assays were conducted similarly to previously described (*9*). Briefly, two 6-mm-diameter-long leaf discs were individually incubated in 50 µM puromycin (Merck) in 24-well-microplates for 1 hour with mild shaking. Afterwards, the two discs per sample were pooled, dried with tissue paper and deep-freeze in liquid nitrogen. Protein extraction and immunoblotting using anti-puromycin antibody (Merck) was conducted as previously described. Relative lane intensities were calculated using the Image Lab software (BioRad).

<u>Omics downstream analysis: Gene ontology and protein network</u>

For Gene Ontology (GO) enrichment, the lists of *A. thaliana* genes were uploaded into ShinyGO software (*54*). Table of annotations was extracted for further analysis. For the generation of protein networks, the lists of *A. thaliana* genes were provided to the STRING database (*55*). For purposes of clarity, network nodes were manually arranged. Clusters were automatically generated using the k-mean clustering tool, and representative GO terms were manually assigned based on the enrichment test automatically conduced within the STRING software. Representation of the protein networks was done with Cytoscape (*56*).

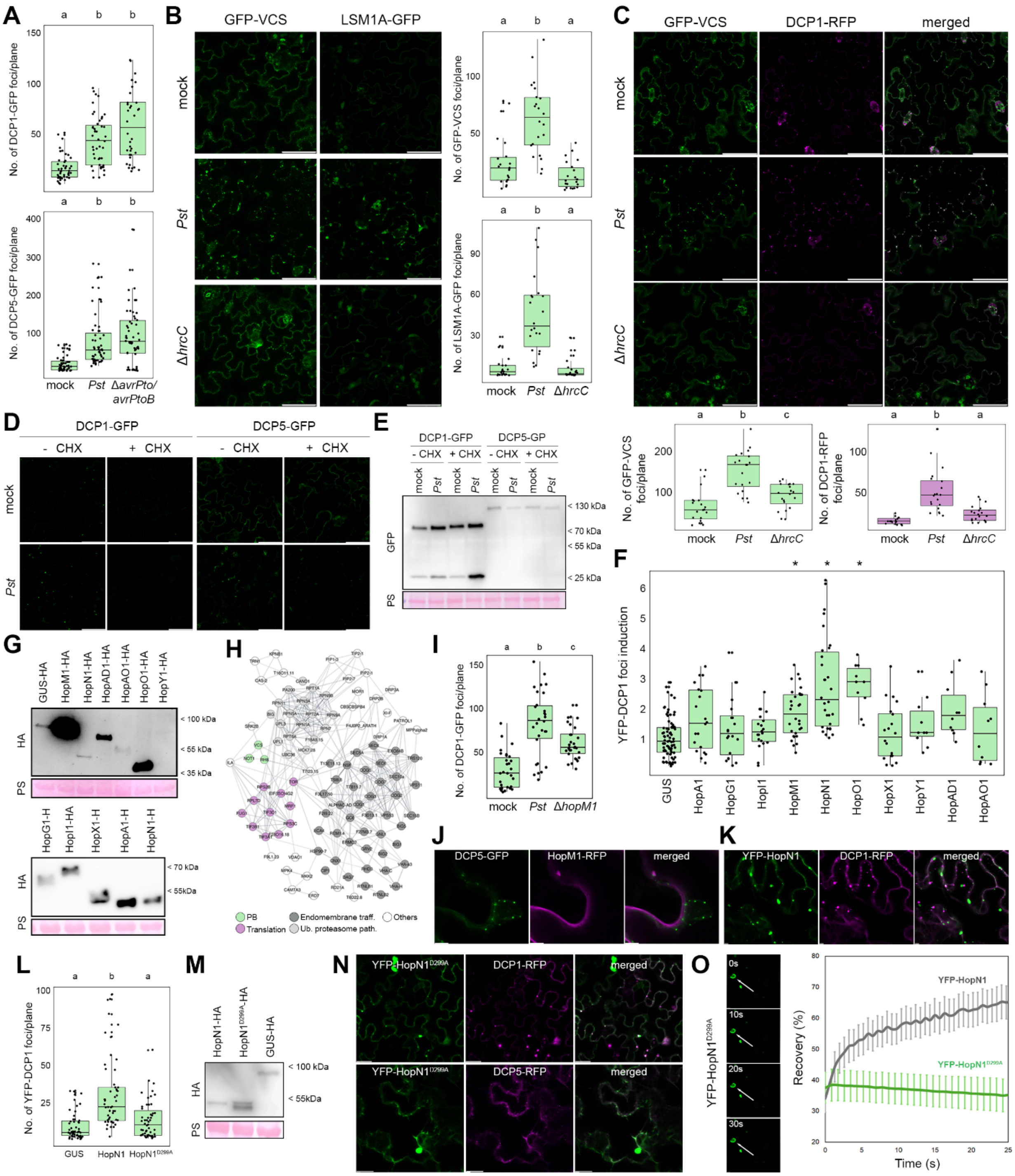


**Fig. S1. *Pseudomonas syringae* induces P-body formation in an effector-dependent manner.**

(**A**-**C**) Confocal microscopy pictures and boxplots representing the number of fluorescent foci per plane observed in epidermal cells of 4-week-old *A. thaliana* leaves expressing DCP1-GFP or DCP5-GFP (A), GFP-VCS or LSM1A-GFP (B), and GFP-VCS and DCP1-RFP (C) infected with *Pst* wild-type and mutant strains or 10 mM $MgCl^2$ mock at 24 hours post mesophyll-infiltration with $10^7$ CFUs/mL. Scale bars depict 50 µm. Data from at least two independent experiments. Different letters indicate statistical groups determined using Student *t*-test (p-value < 0.05). (**D**) Confocal microscopy pictures of epidermal cells of 4-week-old *A. thaliana* leaves mesophyll-infiltrated with 10 mM $MgCl_2$ mock or $10^7$ CFUs/mL *Pst* 24 hours after inoculation and 10 mM $MgCl_2$ control or 100 µM cycloheximide (CHX) 4 hours before imaging. Scale bars depict 50 µm. Representative images from two independent experiments. (**E**) Immunoblot analysis of DCP1-GFP and DCP5-GFP protein levels in 4-week-old A. thaliana leaves mock or Pst-infected and control or CHX-treated from which images were shown in D. The large subunit of the Rubisco visible after Ponceau S (PS) staining serves as loading control. Representative images from two independent experiments. (**F**) Boxplot representing the number of fluorescent foci per plane observed in epidermal cells of 5-week-old *N. benthamiana* leaves transiently expressing YFP-DCP1 and HA-tagged *Pst* effectors or HA-tagged β-glucuronidase (GUS) as negative control at 24 hours post mesophyll. Data normalized to the average of the HA-tagged GUS negative control of each independent experiment. Data from at least two independent experiments. Asterisk indicates statistically significant differences compared to HA-tagged GUS negative control (Student *t*-test p-value < 0.05). (**G**) Immunoblot analysis of HA-tagged *Pst* effectors and GUS control transiently expressed in 5-week-old *N. benthamiana* leaves at 48 hours after inoculation. The large subunit of the Rubisco visible after Ponceau S (PS) staining serves as loading control. (**H**) Protein network of interactors of HopM1 according to a previous publication (*21*). Groups were manually assigned based on representative gene ontology (GO) terms. (**I**) Number of fluorescent foci per plane observed in epidermal cells of 4-week-old *A. thaliana* leaves expressing DCP1-GFP infected with *Pst* wild-type and Δ*hopM1* mutant or 10 mM $MgCl_2$ mock at 24 hours post mesophyll-infiltration with $10^7$ CFUs/mL. Data from two independent experiments. Different letters indicate statistical groups determined using Student *t*-test (p-value < 0.05). (**J-K** and **N**) Confocal microscopy pictures of transiently co-expressed GFP-DCP5 with HopM1-RFP (J); YFP-HopN1 with DCP1-RFP (K); or YFP-$HopN1^{D299A}$ with DCP1-RFP or DCP5-RFP (N) in 5-week-old *N. benthamiana* epidermal cells. Images were taken 30–48 h after infiltration. Scale bars represent 5 µm. (**L**) Number of fluorescent foci per plane observed in epidermal cells of 5-week-old *N. benthamiana* leaves transiently expressing YFP-DCP1 and HA-tagged HopN1, catalytic-dead mutant $HopN1^{D299A}$ or GUS as negative control at 48 hours post mesophyll-infiltration. Data from three independent experiments. Different letters indicate statistical groups determined using Student *t*-test (p-value < 0.05). (**M**) Immunoblot analysis of HA-HopN1, $HopN1^{D299A}$ or GUS control transiently expressed in 5-week-old *N. benthamiana* leaves from the plants whose images were quantified in L. The large subunit of the Rubisco visible after Ponceau S (PS) staining serves as loading control. (**O**) Representative confocal microscopy images and time-course plot of the fluorescence signal recovery of YFP-$HopN1^{D299A}$ foci after photobleaching (green). Datra from YFP-HopN1 from Fig. 1H represented in grey for comparison. Recovery was quantified as the percentage of fluorescence intensity compared to before photobleaching. The data represented is the mean and error bars represent the standard error of the mean (n > 10). The scale bar depicts 5 µm.

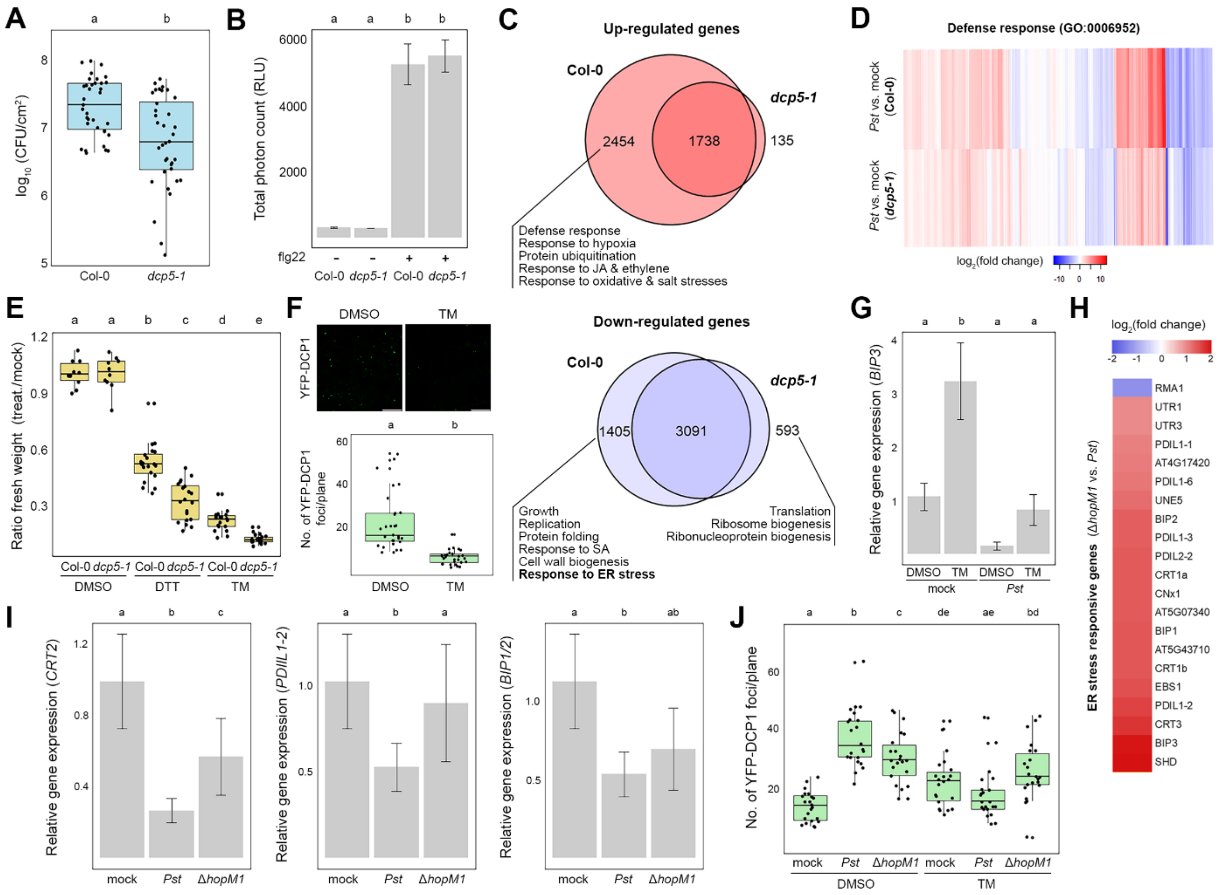


**Fig. S2. P-bodies are positive regulators of susceptibility via an altered ER stress response.**

(**A**) Bacterial population in 4-week-old *A. thaliana* leaves of wild type Col-0 and *dcp5*-1 plants mesophyll-infiltrated with *Pst* solution at $10^4$ CFU/mL three days post-inoculation. Data from three independent experiments. Different letters indicate statistical groups determined using Student *t*-test (p-value < 0.05). (**B**) Total production of reactive oxygen species (ROS) during one hour upon elicitation, or not, with 50 nM flg22 in 4-week-old *dcp5-1* and Col-0 *A. thaliana* plants measured by luminometry and expressed in relative light units (RLUs). Data from two independent experiments. Different letters indicate statistical groups determined using Student *t*-test (p-value < 0.05). (**C**) Venn diagrams depicting the overlap of differentially up- (in red, top) and down- (in blue, bottom) regulated genes in 4-week-old *dcp5-1* and Col-0 plants 24 hours after mesophyll-infiltration with 10 mM $MgCl_2$ mock or $10^7$ CFUs/mL *Pst*. Numbers within the Venn diagrams indicate the number of DEG for each condition ( | $log_2$(fold change) | > 1.5 and adjusted p-value < 0.05). Enriched GO terms (biological process) of each exclusive section are depicted below. JA: jasmonic acid, SA: salicylic acid, and ER: endoplasmic reticulum. (**D**) Heatmap representing the $log_2$(fold change) of defense response genes in transcriptomic analyses depicted in Fig. 2. (**E**) Ratios of relative fresh weight (treatment/mock) of 2-week-old seedlings grown in DMSO mock and ER stress-inducing drugs dithiothreitol (DTT; 1.5 mM) or tunicamycin (TM; 150 nM). Data from two independent experiments. Different letters indicate statistical groups determined using Student *t*-test (p-value < 0.05). (**F**) Confocal microscopy pictures and number of fluorescent foci per plane observed in the root meristem of 7-day-old *A. thaliana* seedlings expressing YFP-DCP1 incubated for 8 hours in 6 µM TM or equivalent DMSO volume-supplemented liquid medium. Data from two independent experiments. Different letters indicate statistical groups determined using Student *t*-test (p-value < 0.05). The scale bar represents 25 µm. (**G**) Relative gene expression of ER stress response gene *BIP3* in 4-week-old *A. thaliana* plants infected with either 10 mM MgCl2 mock or $10^7$ CFUs/mL *Pst* and either 10 µM TM or equivalent DMSO volume 24 and 8 hours prior imaging, respectively. Error bars depict the standard error of the mean. Data from two independent experiments. Different letters indicate statistical groups determined using Student *t*-test (p-value < 0.05). (**H**) Heatmap representing the $log_2$(fold change) of certain ER stress response-related genes upon infection of Δ*hopM1* mutants vs. wild-type *Pst* from a previously published data (*28*). (**I**) Relative gene expression of ER stress response genes *CALRETICULIN 2* (*CRT2*), *PROTEIN DISULFIDE ISOMERASE-LIKE 1-2* (*PDIL1-2*) and *BIP1/2* in 4-week-old *A.*

*thaliana* plants 24 hours after infection with either 10 mM MgCl2 mock or $10^7$ CFUs/mL wild-type *Pst* or single Δ*hopM1* mutant. Error bars depict the standard error of the mean. Data from three independent experiments. Different letters indicate statistical groups determined using Student *t*-test (p-value < 0.05). (**J**) Number of fluorescent foci per plane observed in epidermal cells of 4-week-old *A. thaliana* leaves expressing YFP-DP1 infected with either 10 mM MgCl2 mock or $10^7$ CFUs/mL of wild-type *Pst* or single Δ*hopM1* mutant and either 6 μM TM or equivalent DMSO volume, 24 and 8 hours prior imaging, respectively. Data from two independent experiments. Different letters indicate statistical groups determined using Student *t*-test (p-value < 0.05).

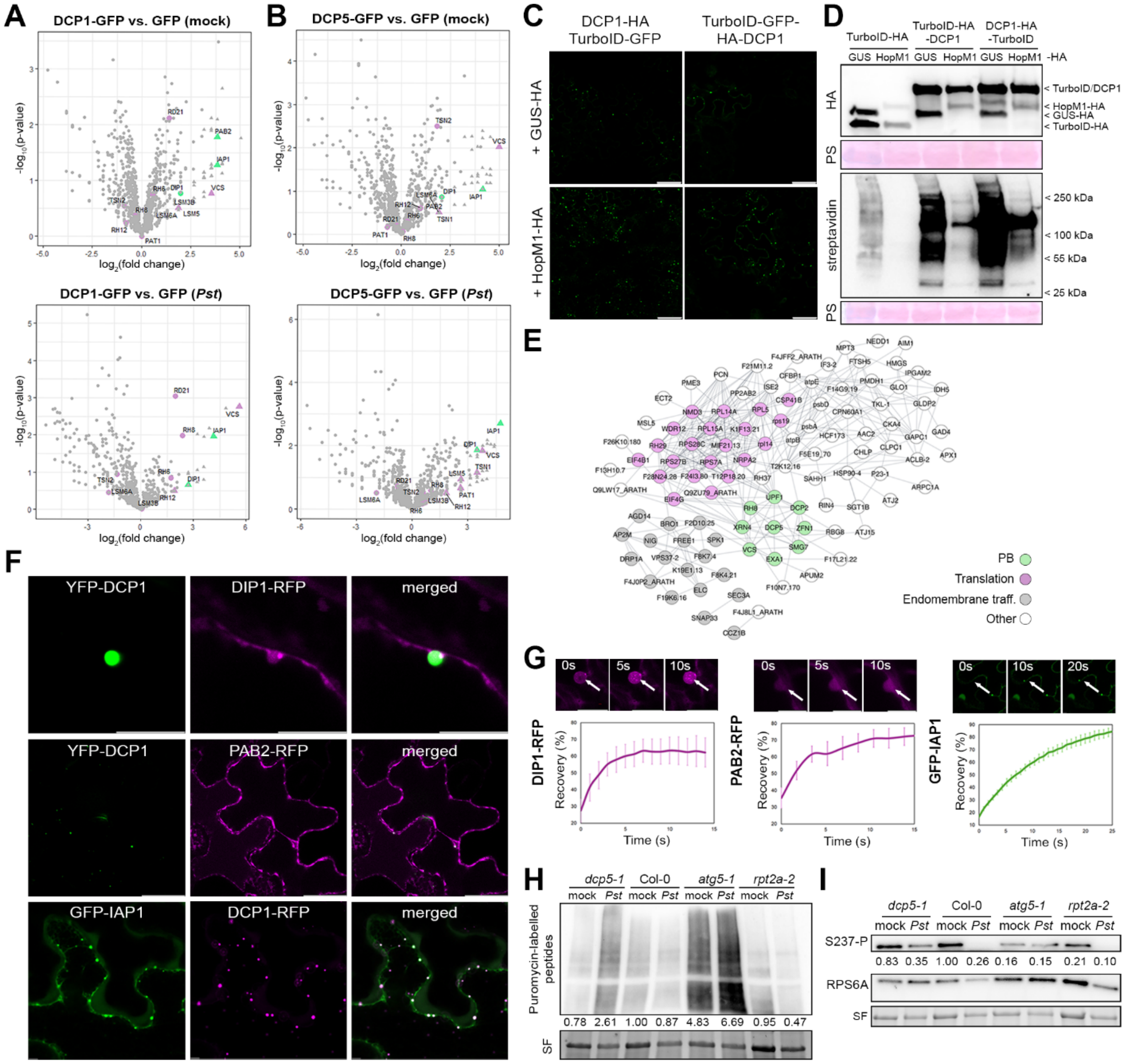


**Fig. S3. P-bodies are required for *Pst*-mediated attenuation of general translation.**

(**A**-**B**) Volcano plots representing the results of the IP-MS/MS proteome analysis of proteins enriched in either DCP1-GFP (A) or DCP5-GFP (B)-expressing plants, compared to GFP control 24 hours upon vacuum-infiltration with either 10 mM $MgCl_2$ mock (left) or $10^7$ CFUs/mL *Pst* infection. Highlighted in purple are known P-body components, and in green, candidates further studied in this work. (**C**) Confocal microscopy pictures of epidermal cells of 5-week-old *N. benthamiana* leaves transiently co-expressing DCP1-HA-TurboID-GFP and TurboID-GFP-HA-DCP1 with either HA-tagged HopM1 or GUS as negative control. Images were taken 48 h after infiltration. The scale bar represents 50 µm. (**D**) Test for expression and biotinylation activity of DCP1-HA-TurboID-GFP, TurboID-GFP-HA-DCP1 and TurboID-GFP-HA control transiently co-expressed with either HA-tagged HopM1 or GUS in 5-week-old *N. benthamiana* leaves 48 hours after infiltration. The HA immunoblot analysis (top) shows the presence of all constructs whereas the streptavidin blot (bottom) shows the biotinylation profile of each construct upon supplementation with 50 µM biotin for 6 hours. The large subunit of the Rubisco visible after Ponceau S (PS) staining serves as loading control. (**E**) Protein network of the biotinylated proteins enriched in 5-week-old *N. benthamiana* leaves transiently expressing DCP1-HA-TurboID-GFP or TurboID-GFP-HA-DCP1 compared to TurboID-GFP-HA negative control 48 hours after infiltration. Groups were manually assigned based on representative gene ontology (GO) terms enriched in different clusters obtained by k-mean clustering in STRING. (**F**) Confocal microscopy pictures of epidermal cells of 5-week-old *N. benthamiana* leaves transiently co-expressing YFP-DCP1 and DIP1-RFP (top), YFP-DCP1 and PAB2-RFP (middle) and GFP-IAP1 and DCP1-RFP (bottom). Images were taken 48 h after infiltration. Scale bars represent 10 µm (top), 50 µm (middle) and 5 µm (bottom). (**G**) Representative confocal microscopy images and time-course plot of the fluorescence signal recovery of DIP1-RFP (left), PAB2-RFP (centre) and GFP-IAP1 (right) foci after photobleaching. Recovery was quantified as the percentage of fluorescence intensity compared to before photobleaching. The data represented is the mean and error bars represent the standard error of the mean ($n > 10$). (**H-I**) Immunoblot analysis of puromycin-labelled peptides (H) or phosphorylation status at Ser237 of RPS6A (I) in 4-week-old Col-0, *dcp5-1*, *atg5-1* and *rpt2a-2 A. thaliana* leaves 24 hours after mesophyll-infiltration with either 10 mM MgCl2 mock or $10^7$ CFUs/mL *Pst*. The large subunit of the Rubisco visible through Stain-Free (SF) imaging revelation serves as loading control. Numbers indicate the lane intensity of the puromycin-labelled peptide immunoblot normalized to each sample loading (H) or the band intensity ratio between the phosphorylation-specific RPS6A immunoblot and its respective phosphorylation-insensitive RPS6A blots (I). Representative results from two independent experiments.

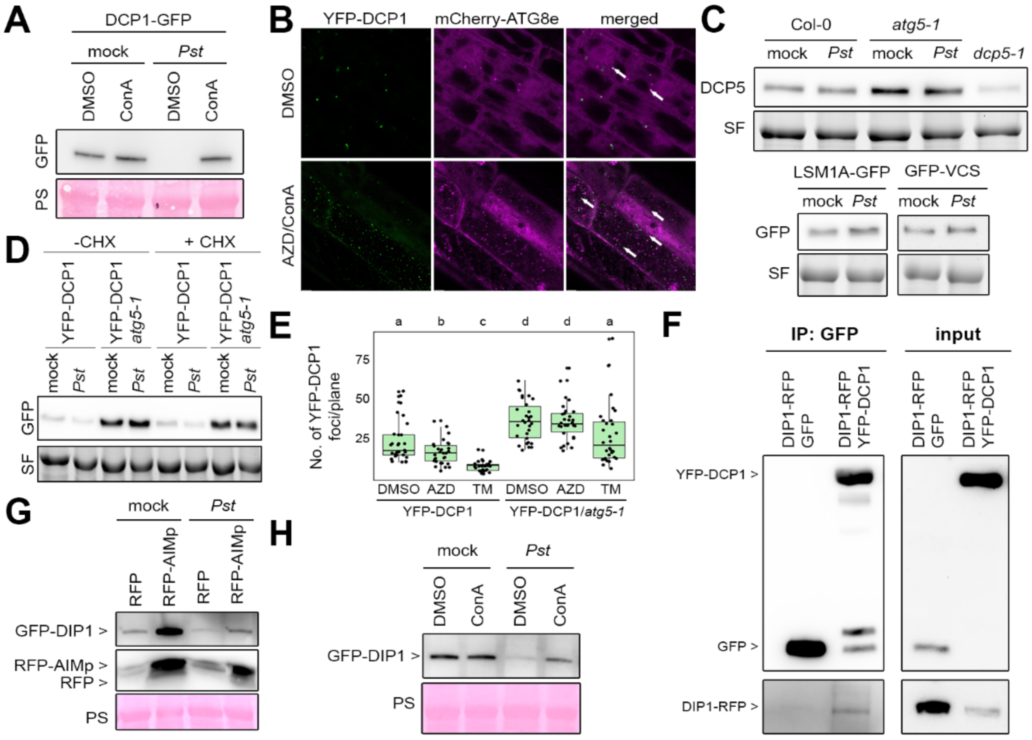


**Fig. S4. *Pst* utilizes autophagy to modulate P-body dynamics.**

(**A**) Immunoblot analysis of DCP1-GFP levels in 4-week-old DCP1-GFP *A. thaliana* plants mesophyll-infiltrated with 10 mM $MgCl_2$ mock or $10^7$ CFUs/mL *Pst* and vacuum-infiltrated with 1 µM concanamycin A (ConA) or equivalent DMSO volume for 24 and 6 hours before harvesting respectively. The large subunit of the Rubisco visible through Ponceau S (PS) staining serves as loading control. Representative result from two independent experiments. (**B**) Confocal microscopy pictures of root meristem of 7-day-old *A. thaliana* seedlings expressing YFP-DCP1 and mCherry-ATG8E treated with 10 µM AZD8055 (AZD) and 1 µM ConA or equivalent DMSO volume for 6 hours prior to imaging. White arrows highlight co-localization of YFP-DCP1 and mCherry-ATG8E foci. Scale bars represent 5 µm. (**C**) Immunoblot analysis of DCP5 protein levels in 4-week-old Col-0, *atg5-1* and *dcp5-1* (top) or GFP-tagged protein levels in 4-week-old LSM1A-GFP and GFP-VCS (bottom) *A. thaliana* plants mesophyll-infiltrated with 10 mM $MgCl_2$ mock or $10^7$ CFUs/mL *Pst* 24 hours before harvesting. The large subunit of the Rubisco visible through Stain-Free (SF) imaging revelation serves as loading control. Representative results from two independent experiments. (**D**) Immunoblot analysis of YFP-DCP1 levels in 4-week-old *A. thaliana* leaves expressing YFP-DCP1 in the Col-0 and *atg5-1* background mesophyll-infiltrated with 10 mM $MgCl_2$ mock or $10^7$ CFUs/mL *Pst* and vacuum-infiltrated with 20 µM CHX or equivalent water volume 24 and 4 hours before harvesting, respectively. The large subunit of the Rubisco visible through Stain-Free (SF) imaging revelation serves as loading control. (**E**) Quantification of fluorescent foci per plane observed in the root meristem of 7-day-old *A. thaliana* seedlings expressing YFP-DCP1 in Col-0 and *atg5-1* background incubated for 8 hours in 6 µM TM, 10 µM AZD8055 or equivalent DMSO volume-supplemented liquid medium. Data from two independent experiments. Different letters indicate statistical groups determined using Student *t*-test (p-value < 0.05). (**F**) Coimmunoprecipitation of YFP-DCP1 or GFP with DIP1-RFP transiently expressed in 5-week-old *N. benthamiana* leaves 48 hours after inoculation. Total proteins (input) were subjected to immunoprecipitation (IP) with GFP-Trap beads, followed by immunoblot analysis using either anti-GFP and -RFP antibodies. Representative result from two independent experiments. (**G**) Immunoblot analysis of GFP- and RFP-tagged protein levels in 5-week-old *N. benthamiana* leaves transiently co-expressing GFP-DIP1 with either RFP-AIM peptide (AIMp) or RFP for 48 hours, and mesophyll-infiltrated with either 10 mM $MgCl_2$ mock or $10^7$ CFUs/mL Δ*hopQ1 Pst* for 8 hours. The large subunit of the Rubisco visible through Ponceau S (PS) staining serves as loading control. Representative result from two independent experiments. (**H**) Immunoblot analysis of GFP-tagged protein levels in 5-week-old *N. benthamiana* leaves transiently expressing GFP-DIP1 for 48 hours, and mesophyll-infiltrated with either 10 mM $MgCl_2$ mock or $10^7$ CFUs/mL Δ*hopQ1 Pst* and 1 μM ConA or equivalent DMSO volume 12 and 6 hours respectively. The large subunit of the Rubisco visible through Ponceau S (PS) staining serves as loading control. Representative result from two independent experiments.

**Table S1. Results from transcriptomic analysis (Col-0 and *dcp5-1* +/- *Pst*) - *Arabidopsis thaliana***

**Table S2. Results from proteomics analysis (IP-MS/MS: GFP, DCP1-GFP and DCP5-GFP +/- *Pst*) - *Arabidopsis thaliana***

**Table S3. Results from proteomics analysis (proximity labelling: GP-TurboID/GFP-TurboID-DCP1/DCP1-TurboID-GP + GUS-HA/HopM1-HA) - *N. benthamiana***

**Table S4. List of *Arabidopsis thaliana* lines used in this study**

**Table S5. List of *Pseudomonas syringae* pathovar tomato (*Pst*) strains used in this study**

**Table S6. List of plasmids used in this study**

**Table S7. List of primers used in this study**

**Table S8. List of antibodies used in this study**
